# Platelets from early-stage Alzheimer patients show enhanced amyloid binding, an elevated open canalicular system and sex-specific differences in their activation profile

**DOI:** 10.1101/2025.07.23.666322

**Authors:** Lili Donner, Julia Christl, Milenko Kujovic, Tillmann Supprian, Margitta Elvers

**Author notes:** **Correspondence:** Margitta Elvers, PhD.

## Abstract

**Introduction:** Alzheimer’s disease (AD) is associated with neurodegeneration and dementia. The clinical parameters include the deposition of amyloid-ß into senile plaques in the brain parenchyma and in cerebral vessels known as cerebral amyloid angiopathy (CAA). Currently, ß-amyloid-antibodies are emerging as possible therapy for AD. Several biomarkers, such as ß-Amyloid and tau-protein have gained significant value for diagnosing early AD. However, these biomarkers require cerebrospinal fluid. Blood tests for screening of AD are urgently needed.

**Methods:** Patients diagnosed with early AD were analyzed for extracellular amyloid-ß binding to platelets, platelet morphology and platelet activation and compared to age-matched controls.

**Results:** Beside unaltered platelet number and size, we detected increased binding of amyloid-ß to activated platelets isolated from AD patients. Electron microscopy revealed an altered platelet morphology in AD patients including the number of dense granules and the area of the open canalicular system (OCS) as compared to controls. While only minor differences in platelet activation were detected between patients and controls, a significant reduction of integrin αIIbβ3 (fibrinogen receptor) activation was evident in platelets from female compared to male AD patients as determined by flow cytometry.

**Conclusion:** The here presented results emphasize the importance to increase our understanding how platelets contribute to AD pathology in patients in a sex-specific manner. Furthermore, platelet parameters might serve as an ideal biomarker for a first prognosis of AD because platelets can be easily accessed by blood samples. These parameters might include a sex-specific platelet activation profile, the capability to bind Aß to the platelet surface and the dimension of the OCS by electron microscopy.

## 1 Introduction

Alzheimer’s disease (AD) is a neurodegenerative disease with progressive decline in cognitive function (Clarfield, 2003). Patients suffering from AD show a specific neuropathological profile: The deposition of extracellular amyloid-ß (Aß) into senile plaques, the formation of intracellular neurofibrillary tangles (NFTs) that arise from hyperphosphorylated tau proteins (Long and Holtzman, 2019). Furthermore, 80% of AD patients develop cerebral amyloid angiopathy that includes the accumulation and aggregation of Aß in cerebral vessels (Selkoe, 2011, Thal et al., 2008). Beside the accumulation of extracellular Aß and intracellular tau, a pathophysiological cascade of neuroinflammation, blood-brain-barrier dysfunction (Thal et al., 2008) leads to neuronal loss and cognitive decline in AD patients. To date, around 57 million people worldwide suffer from dementia (2022). However, with increasing age of the general population and emerging effective therapeutic strategies, there is a strong need for well-established biomarkers and effective therapeutic options that are still lacking today.

Platelets are the smallest blood cells and major regulators of hemostasis and thrombosis, but are also involved in processes of acute and chronic inflammation in disease pathology (Koupenova et al., 2018). It is well known that platelet dysfunction is associated with several neurodegenerative diseases, including Alzheimer’s and Parkinson’s disease (Leiter and Walker, 2020, Beura et al., 2022). Platelets are a source of Aβ peptides and amyloid precursor protein (APP) in the circulation and release different Aß peptides upon activation (Leiter and Walker, 2020, Wolska et al., 2023, Gowert et al., 2014, Li et al., 1998, Chen et al., 1995). Furthermore, platelets are affected by AD with alterations in their activation profile and their adhesion properties (Gowert et al., 2014, Jarre et al., 2014, Sevush et al., 1998, Stellos et al., 2010). In aged transgenic mice modeling Alzheimer’s disease (APP23) with parenchymal plaques and CAA have pre-activated platelets in the circulation and adhere to vascular amyloid-β deposits, leading to cerebral vessel occlusion (Jarre et al., 2014, Gowert et al., 2014, 2022). Beside alterations in platelet physiology, their dysfunction is coupled to different pathological hallmarks in AD such as impairment of mitochondria (Swerdlow et al., 2017), abnormalities in tau production (Forlenza et al., 2011) and alterations in neurotransmitters (Kumar et al., 1995) that affects AD progression.

In our previous studies with an experimental model of AD, the APP23 mice, we identified platelets to convert soluble Aβ40 peptides into fibrillary Aβ aggregates *in vitro* (Gowert et al., 2014, Donner et al., 2020, Donner et al., 2018, Donner et al., 2021, Donner et al., 2016). Furthermore, we were able to provide strong evidence for two direct binding partners of Aβ40 at the platelet surface, the fibrinogen receptor integrin αIIbβ3 and the major collagen receptor glycoprotein VI (GPVI) (Donner et al., 2016, Donner et al., 2020). Targeting these platelet receptors by blocking with an antibody or genetic deletion resulted in a significant reduction of Aβ aggregation *in vitro* (Donner et al., 2016, Donner et al., 2020). Next, we identified adenosine diphosphate (ADP) as major trigger of Aß aggregation at the platelet surface. In detail, we found that ADP plays a role in platelet-mediated Aβ aggregation, as inhibition of the ADP receptor P2Y_12_ prevented Aβ aggregation *in vitro* (Donner et al., 2016). In vivo, treatment of APP23 mice with the antithrombotic drug clopidogrel, a P2Y_12_ antagonist for three months resulted in a significant reduction of CAA in aged APP23 mice (Donner et al., 2016).

To date, different clinical studies exist providing evidence for alterations of platelet parameters in patients with AD. In a cross-sectional study, Sevush and colleagues claim that platelet activation contributes to the pathogenesis of AD. Their results indicated that platelets are a major source of Aß in blood because they found elevated circulating platelet aggregation, elevated P-selectin exposure and increased leukocyte-platelet aggregates (Sevush et al., 1998) 1998). Stellos and colleagues found elevated P-selectin and integrin aIIbb3 at the platelet surface suggesting that platelet activation could serve as a prognostic biomarker for the rate of cognitive decline in AD patients (Stellos et al., 2010). Different researches detected elevated or declined platelet proteins that could be a biomarker for AD patients (Yu et al., 2022, Yu et al., 2021). In addition, alterations in APP ratio of platelets, differences in secretase activity and/or fibrinogen deposition in CAA positive vessels might be a promising biomarker for AD (Di Luca et al., 1998, Fu et al., 2023, Hultman et al., 2013, Johnston et al., 2008, Zainaghi et al., 2012).

In this study, we examined platelet morphology and activation in AD patients at an early stage of the disease. While we detected no differences in platelet count or size, we found an increased binding of soluble Aß, an elevated number of dense granules and an increased area of the open canalicular system (OCS) using platelets from AD patients. With regard to platelet activation, we detected no major differences between AD patients and age-matched healthy controls. However, the analysis of sex-specific platelet parameters revealed significantly reduced platelet activation in female versus male AD patients.

## 2 Materials and Methods

### 2.1 Subjects

A total of 46 patients with AD and 17 healthy elderly controls were included in the study. The diagnosis of Alzheimer’s disease (AD) was based on the criteria of the National Institute of Neurological Disorders and Stroke–Alzheimer Disease and Related Disorders (NINCDS– ADRDA). The clinical severity of cognitive impairment was assessed by the CERAD plus neuropsychological battery.

Age-matched controls from the local blood bank were included in the study.

Experiments with human blood were reviewed and approved by the Ethics Committee of the Heinrich-Heine-University, who approved the collection and analysis of the tissue. Subjects provided informed consent prior to their participation in the study (patients’ consent): Permitted ethical votes; study number 4845R, ID 2014-102828; ID 2018-140KFogU). The study was conducted in accordance with Declaration of Helsinki principles and the International Council for Harmonization Guidelines on Good Clinical Practice.

### 2.2 Flow cytometry

Citrated whole blood was used and diluted 1:10 in human Tyrode’s buffer (137 mM NaCl, 2.8 mM KCl, 12 mM NaHCO_3_, 0.4 mM NaH_2_PO_4_ and 5.5 mM glucose, pH 6.5). Blood samples were mixed with antibodies and stimulated with indicated agonists at RT for 15 min in the dark. The addition of DPBS was used to stop the reaction and the samples were analysed on a FACSCalibur flow cytometer (BD Biosciences).

Briefly, two-color analysis of human platelet activation was performed using fluorophore-labeled antibodies for P-selectin expression (Emfret Analytics, Wug.E9-FITC, #M130-1, and BD Biosciences, anti-human CD62P-PE, #348107) and the active form of α_IIb_β_3_ integrin (Emfret Analytics, JON/A-PE, #M023-2, and BD Biosciences anti-human PAC-1-FITC, #340507) as described recently (Wagenhäuser et al., 2024). For the analysis of surface expression of different glycoproteins as well as activation-dependent upregulation of integrin β_3_ expression, blood samples were mixed with specific antibodies (Integrin α_5_/CD49e, Tap.A12-FITC and anti-human CD61-PE, #555754, BD Biosciences) and incubated for 15 min at RT in the dark.

For determination of Aß binding to platelets, we used a FITC-labeled Aß antibody that detects APP and Aß at the surface of platelets.

### 2.3 Isolation of human platelets

Platelet count and mean platelet volume (MPV) in blood from AD patients and healthy controls were analyzed using a hematology analyzer (Sysmex KX-21N, Norderstedt, Germany). For platelet preparation, blood was centrifuged at 231 *g* for 10 min. Thereafter, the upper phase, i.e. the platelet-rich plasma (PRP), was carefully transferred into Dulbecco’s phosphate buffered saline (DPBS, pH 6.5) containing apyrase (2.5 U/mL) (Sigma-Aldrich, #A7646) and Prostaglandin E1 (PGE_1_, 1 µM) (Sigma-Aldrich, #P5515). The DPBS PRP mixture was centrifuged at 1,000 *g* for 6 min without brakes to induce the formation of a platelet pellet. The pellet was resuspended in Tyrode’s buffer (137 mM NaCl, 2.8 mM KCl, 12 mM NaHCO_3_, 0.4 mM NaH_2_PO_4_ and 5.5 mM glucose, pH 6.5). The cell count was determined using a hematology analyzer (Sysmex KX-21N, Norderstedt, Germany) and adjusted for the following experiment.

### 2.4 Transmission electron microscopy

Non-stimulated platelets were isolated and fixed in Karnovsky’s solution for 1 h at room temperature and stored at 4°C as described recently. Briefly, for electron microscopic studies, cell pellets were embedded in agarose at 37°C, coagulated, cut in small blocks, fixed in Karnovsky’s solutions, postfixed in osmium tetroxide, and embedded in glycid ether and cut using an ultramicrotome (Ultracut Reichert, Vienna, Austria). Ultrathin sections (30 nm) were mounted on copper grids and analyzed using a Zeiss LIBRA 120 transmission electron microscope (Carl Zeiss, Oberkochen, Germany) (Gowert et al., 2014).

### 2.5 Statistical analysis

All statistical analyses were performed with GraphPad Prism (Prism 9; Graph Pad Software, Inc.). Unpaired t-test or one-way ANOVA, for two groups or three groups. The results were presented as the mean with each individual data point or in bar graph ± SEM. A p value <0.05 was considered significant (*p < 0.05, **p < 0.01).

## 3 Results

### 3.1 Normal platelet count and size and unaltered glycoprotein expression but increased Aß binding to platelets from AD patients

For this study, platelets from male and female AD patients were analyzed and compared to age-matched controls. As shown in Suppl.-figure 1A, all subjects included in this study were >70 years of age (Suppl.-figure 1A). Platelet count and size (mean platelet volume, MPV) in blood from AD patients and healthy controls were analyzed using a hematology analyzer (Sysmex KX-21N, Norderstedt, Germany). As shown in figure 1A-B, no differences have been observed between AD patients and age-matched controls (figure 1A-B). A sex-specific analysis revealed again no differences in platelet account and MPV (Supplemental figure 1B-C) suggesting that there are no differences, neither between AD patients and healthy controls nor between female and male AD patients.

**Figure 1.**
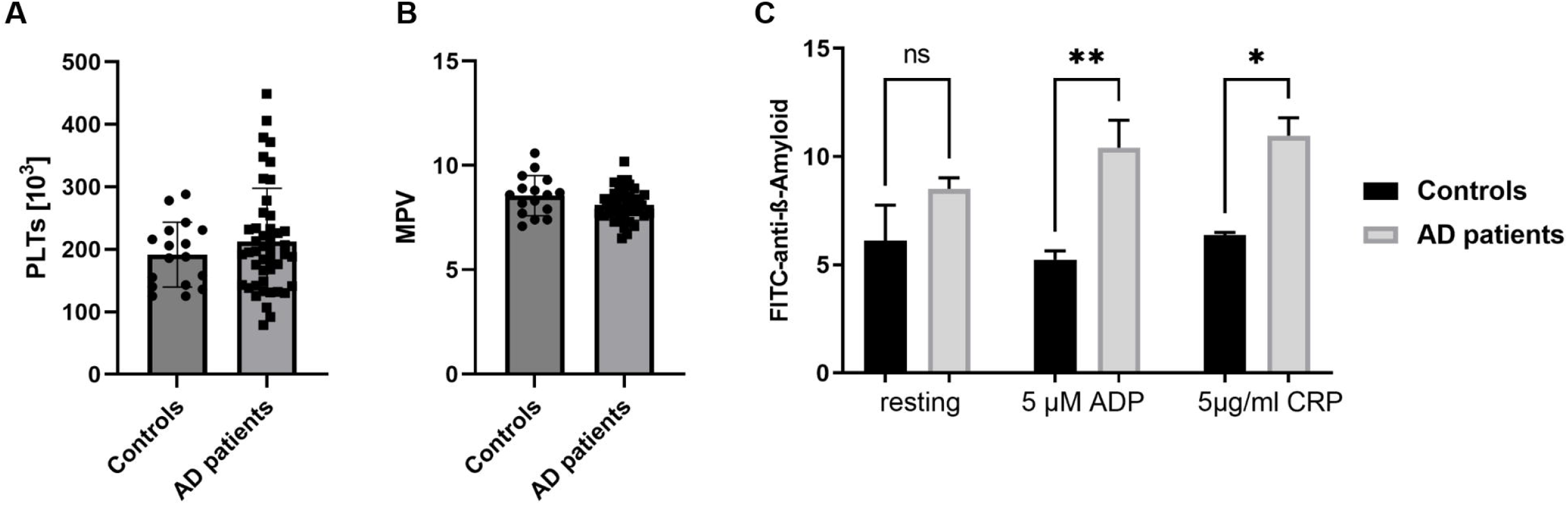
Normal platelet count and size but increased Aß binding to platelets from AD patients. (A) Platelet count and (B) MPV (platelet size) in AD patients and age-matched controls (n = controls 17, patients 46). (C) Binding of Aß to platelets under resting (non-stimulating) and activated conditions was determined by flow cytometry using a FITC-labeled antibody against Aß and APP. Different agonists were used to induce the activation of different signaling pathways in platelets. Results are represented as MFI. Bar graphs indicate mean values ± SEM. Statistical analyses were performed using a multiple unpaired t-test. *p < 0.05; **p < 0.01. ADP, adenosinediphosphate, CRP, collagen-related peptide, MPV, mean platelet volume, MFI, mean fluorescence intensity.

Next, we analyzed the binding of Aß to platelets under resting (non-stimulating) and activated conditions using ADP to activate the ADP receptors P2Y1 and P2Y12 and CRP to activate the major collagen receptor at the platelet surface GPVI. As shown in figure 1C, no differences have been observed under resting conditions. However, when we activated platelets, we detected elevated binding of Aß to platelets from AD patients compared to healthy subjects.

To analyze, if elevated glycoprotein expression is responsible for increased binding of Aß to the surface of platelets from AD patients, we determined the expression of glycoproteins using flow cytometry. However, no differences were detected. The exposure of GPVI, a5-integrin, GPIb and CD61 at the surface of resting platelets was unaltered between AD patients and age-matched healthy controls (figure 2A-D). Also, sex-specific analysis of glycoprotein expression revealed no differences between male and female AD platelets (supplemental figure 2A-D).

**Figure 2.**
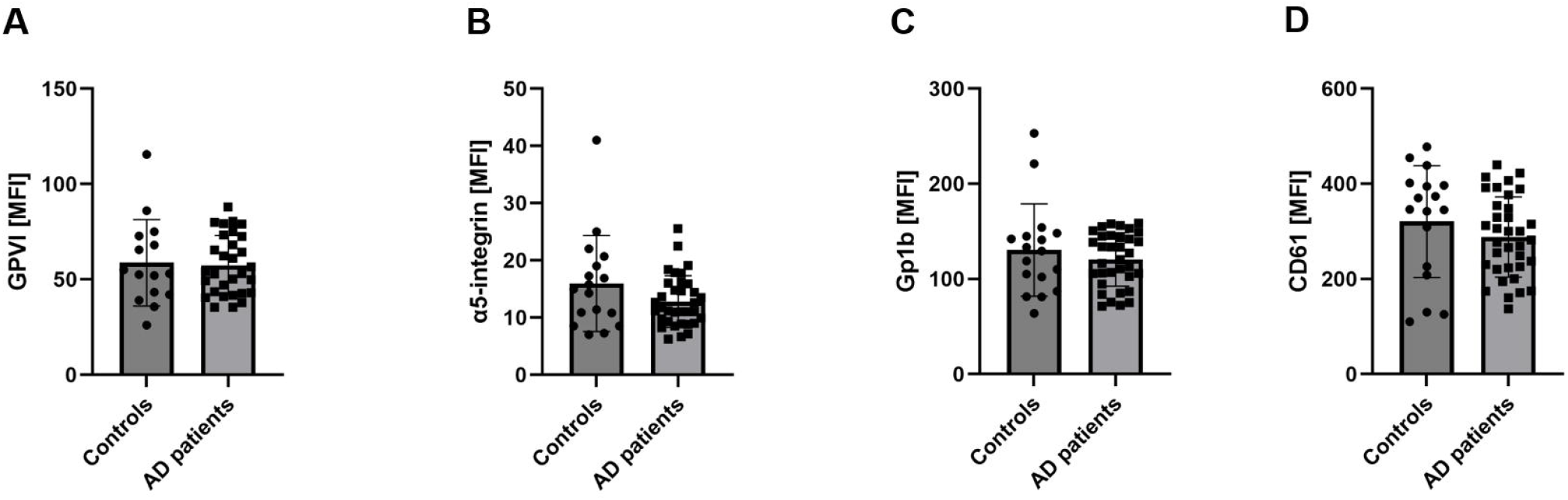
Unaltered glycoprotein expression at the surface of platelets from AD patients. Glycoprotein exposure at the platelet surface was determined by flow cytometry using different antibodies against (A) GPVI, (B) a5-integrin, (C) GPIb and (D) CD61 (subunit of integrin aIIbb3. Results are represented as MFI. Bar graphs indicate mean values ± SEM. Statistical analyses were performed using an unpaired t-test. N = 17 (controls), 35 (patients). MFI, mean fluorescence intensity.

### 3.2 Transmission electron microscopy revealed differences in the number of granules in platelets from AD patients

To analyze platelet morphology in more detail, we performed transmission electron microscopy (TEM). First, we analyzed the number of dense and alpha granules in platelets from AD patients and compared the results to healthy subjects. As shown in figure 3, significantly less platelets from AD patients displayed no dense granules while no differences were detected with platelets containing 1 and 2 dense granules per section (figure 3A, C). Thus, platelets from AD patients contain more dense granules than platelets from healthy subjects. Furthermore, we analyzed the number of alpha granules. No differences were observed between platelets from AD patients and healthy subjects. Most of the platelets from both groups displayed more than 5 alpha granules per platelet as analyzed by TEM (figure 3B, C).

**Figure 3.**
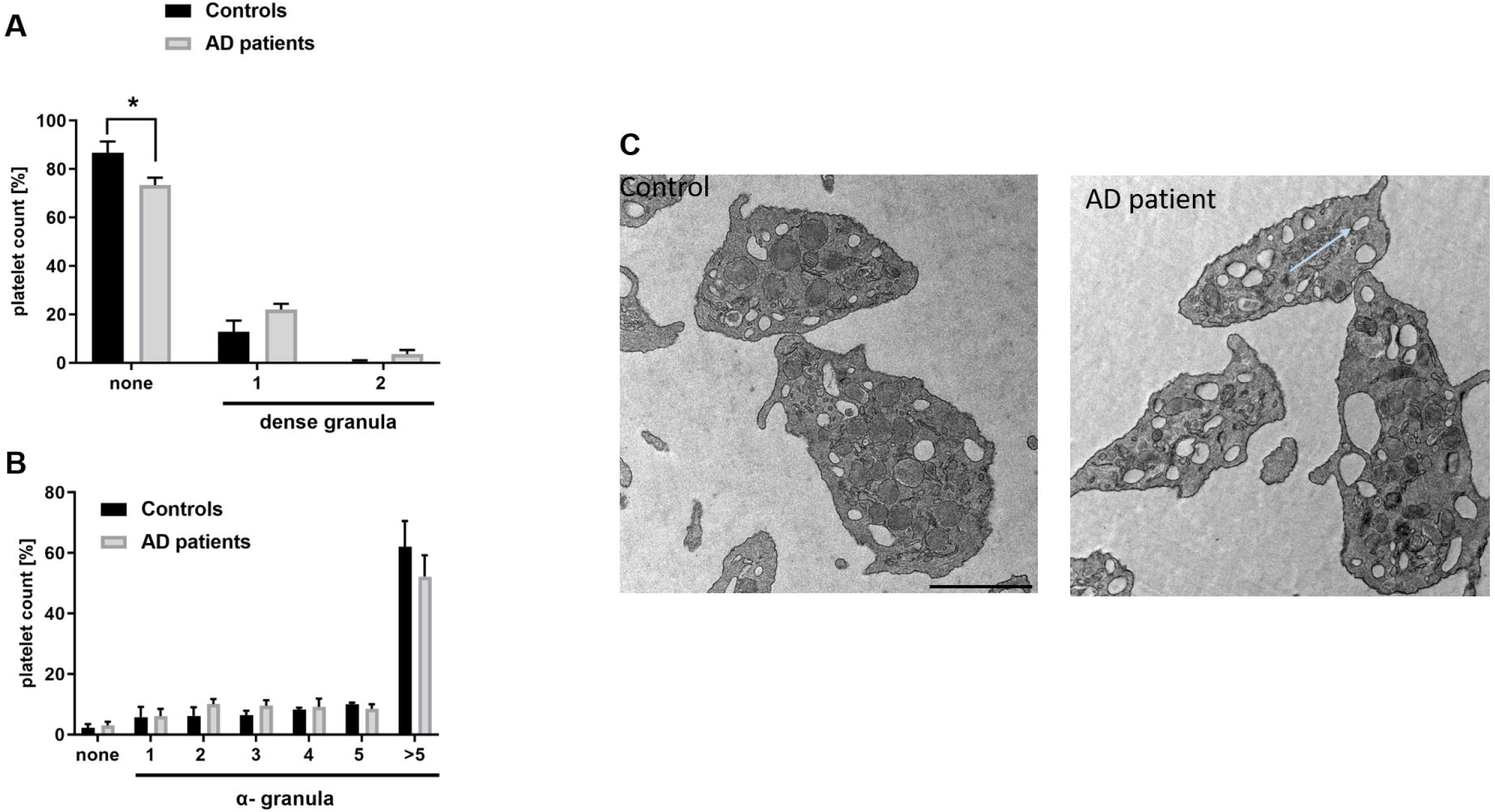
Different number of granules in platelets from AD patients. Analysis of the number of (A) dense and (B) alpha granules in platelets from AD patients and healthy subjects by transmission electron microscopy. (C) Representative images are shown. Scale bar 1 µm, Statistical analyses were performed using an unpaired t-test. N=4-5.

### 3.3 Significantly increased open canalicular system in platelets from AD patients

In a second approach, the open canalicular system of platelets was investigated. To this end, we detected a significantly enhanced area of the OCS in platelets from AD patients when compared to platelets from healthy controls (figure 4A-B). TEM images revealed large areas of the OCS in AD platelets compared to controls (figure 4B). Thus, a detailed analysis of the morphology including the number of granules and the size of the OCS of platelets from AD patients might be promising for the establishment of a prognostic biomarker for AD.

**Figure 4.**
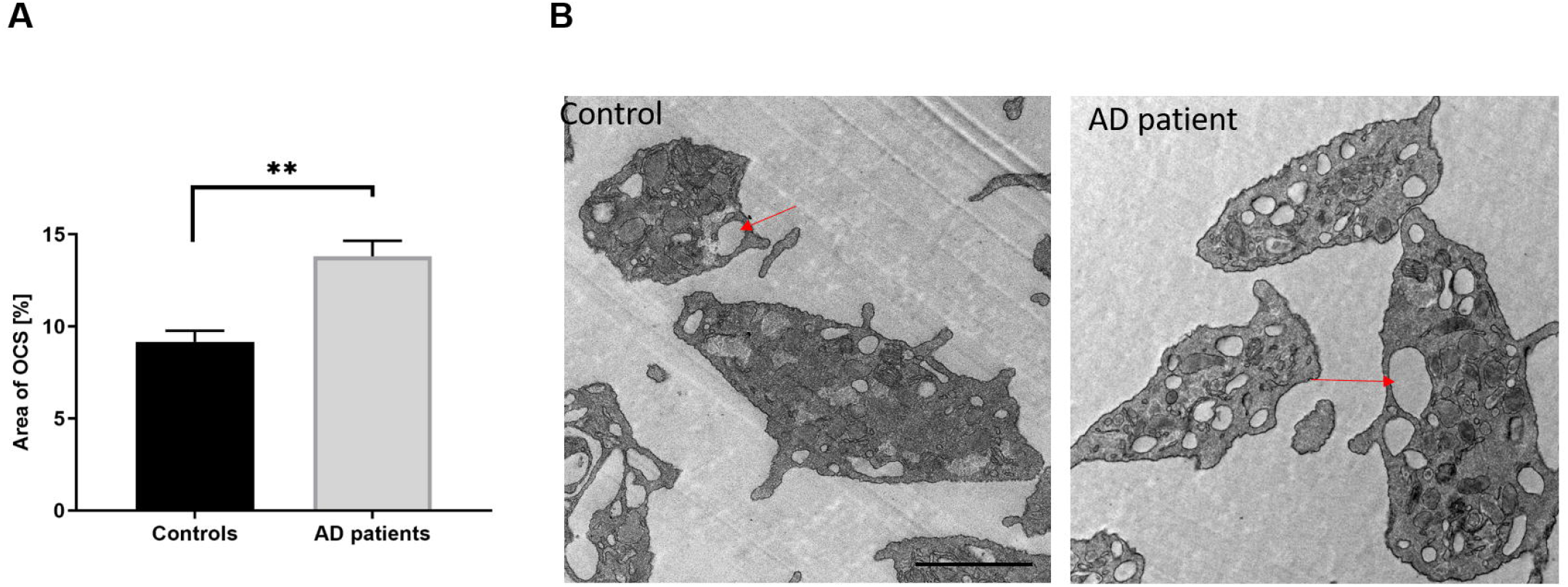
Significantly increased open canalicular system in platelets from AD patients. The area of the open canalicular system of platelets has been analyzed by transmission electron microscopy. (A) Bar graph depicts mean values ± SEM. (B) Representative images from transmission electron microcopy. Scale bar 1 µm. Statistical analyses were performed using an unpaired t-test. N = 3-5.

### 3.4 No major differences in the activation profile of platelets from AD patients

To compare the activation profile of platelets from AD patients and age-matched controls, we determined active integrin aIIbb3 and P-selectin exposure at the platelet surface of both groups. Thereby, the determination of P-selectin exposure serves as marker for degranulation of alpha granules while active integrin aIIbb3 is able to bind the ligand fibrinogen that bridges platelets and is therefore important for platelet aggregation and thrombus formation. under non-stimulating (resting) conditions, no differences in integrin activation or P-selectin exposure were observed between groups (figure 5A-B). next, we analyzed platelet activation following stimulation of platelets by different agonists that activate different signaling pathways in platelets. While low dose ADP, Aß, CRP and PMA induce integrin activation (figure 5C) and P-selectin exposure at the platelet surface (figure 5D), no differences between groups have been detected. However, a significant decrease in P-selectin exposure was observed when we stimulated platelets with 10 µM ADP (figure 5D). Taken together, we did not detect major differences in platelet activation between AD patients and healthy controls (figure 5A-D).

**Figure 5.**
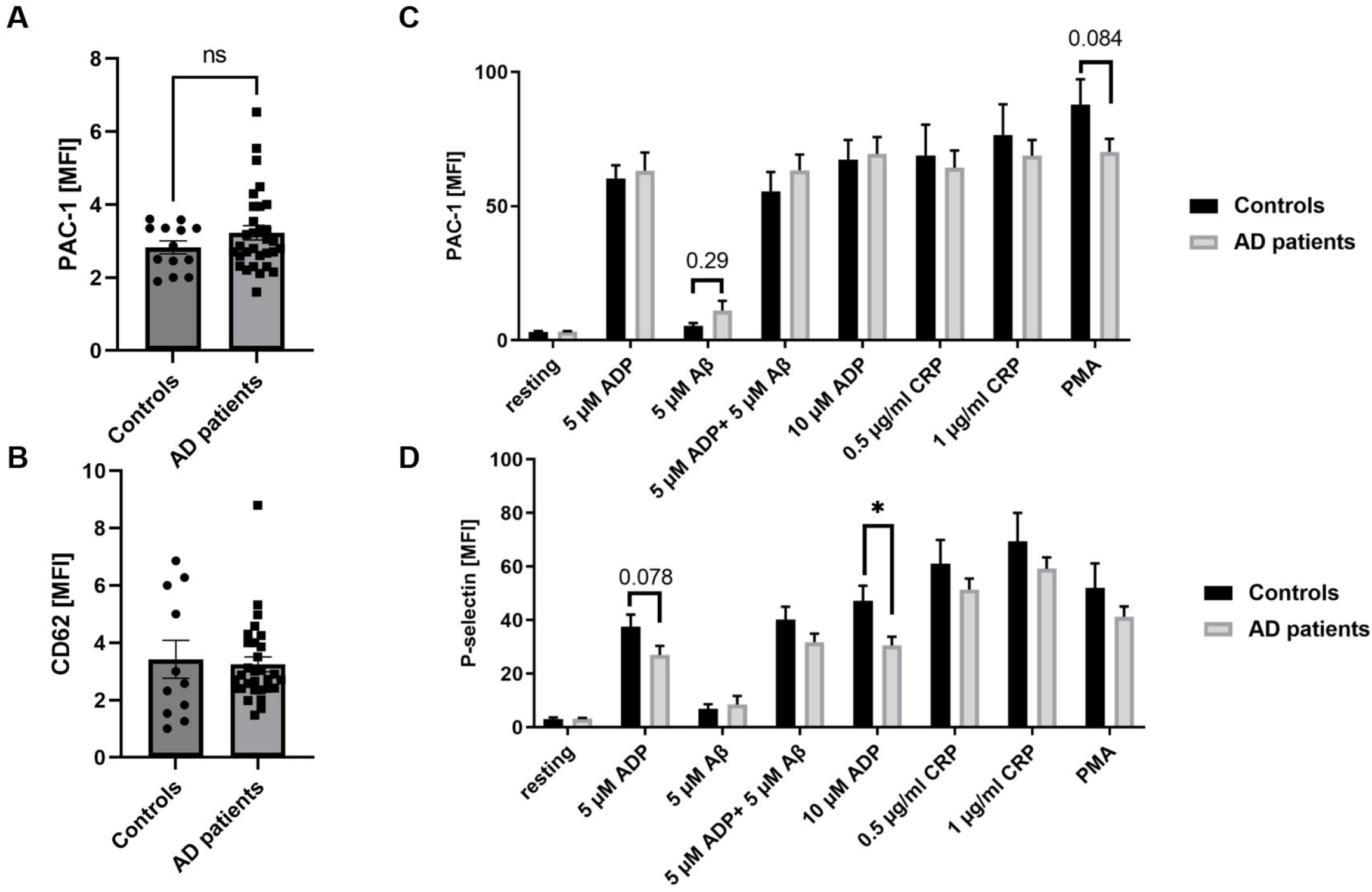
Platelets from AD patients show no major differences in platelet integrin activation and degranulation. (A) Active integrin (PAC-1 binding to integrin aIIbb3) and (B) platelet degranulation (P-selectin-PE) under resting conditions were determined by flow cytometry using whole blood from AD patients and age-matched controls (n = 11 (controls) and 30 (patients). (C-D) Active integrin and P-selectin exposure under activated conditions using different agonists to stimulate different signaling pathways in platelets. Data are represented as MFI. Bar graphs indicate mean values ± SEM. Statistical analyses were performed using a multiple unpaired t-test. *p < 0.05. ADP, adenosinediphosphate, CRP, collagen-related peptide, Aß, amyloid-beta, PMA, Phorbol-myristate-acetate, MFI, mean fluorescence intensity. N = 11 (controls), 30 (patients).

### 3.5 Sex-specific alterations provide evidence for reduced integrin activation of female platelets isolated from AD patients

Next, we investigated if there are any sex-specific differences occurring in platelets from AD patients. As shown in figure 6A, significantly reduced integrin activation was detected when we stimulated the collagen receptor GPVI at the surface of female compared to male platelets from AD patients. Interestingly, under non-stimulating (resting) conditions, significant differences could be also observed. In detail, platelets from female AD patients show less integrin activation under resting conditions (figure 6A). In contrast, no differences have been observed with regard to P-selectin exposure. Thus, it is tempting to speculate that sex-specific analysis of the platelet activation profile might be important to investigate platelet alterations and to establish a prognostic marker for AD.

**Figure 6.**
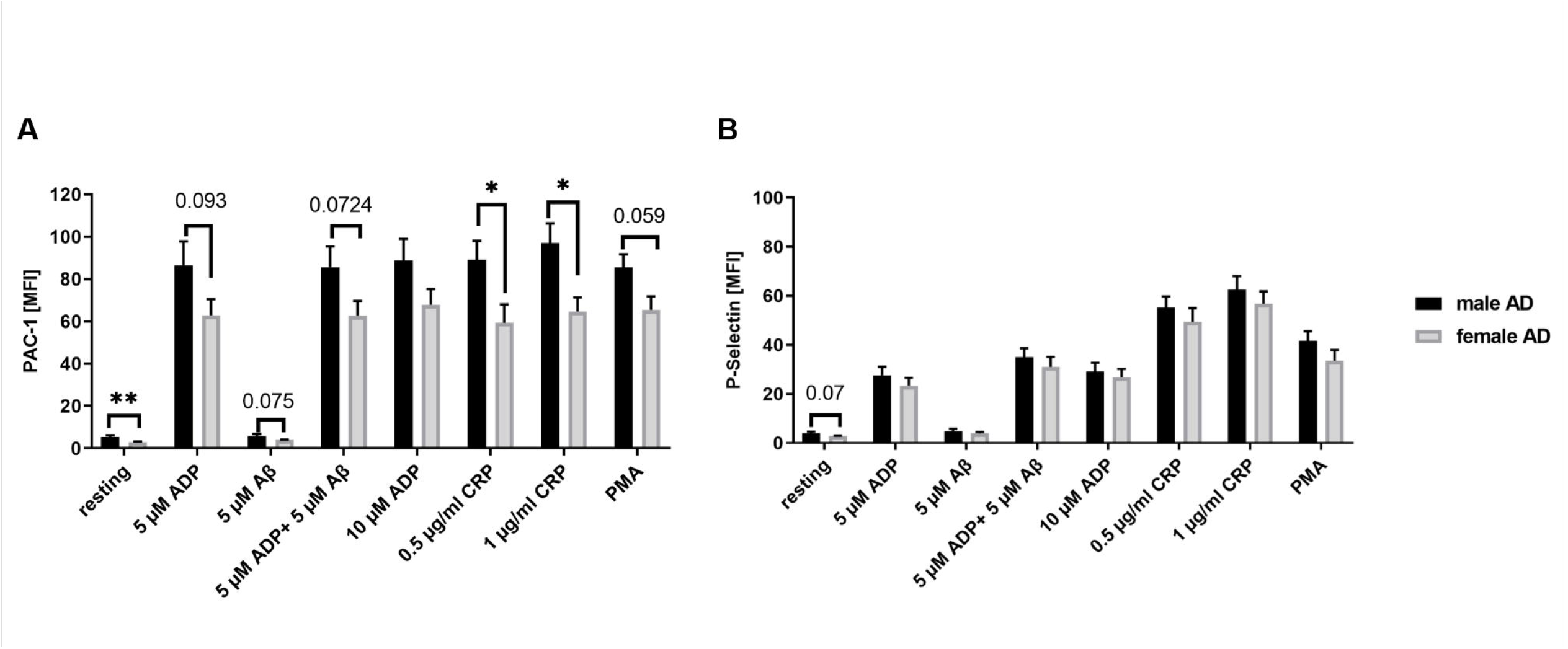
Sex-specific analysis of platelet activation revealed a reduced activation profile in female patients. (A) Active integrin aIIbb3 (PAC-1 binding) and (B) P-selectin exposure as marker for degranulation were determined by flow cytometry. Data are represented as MFI. Bar graphs indicate mean values ± SEM. Statistical analyses were performed using a multiple unpaired t-test. *p < 0.05. ADP, adenosinediphosphate, CRP, collagen-related peptide, Aß, amyloid-beta, PMA, Phorbol-myristate-acetate, MFI, mean fluorescence intensity. N = 11 (controls), 30 (patients).

## 4 Discussion

AD is a multifactorial disorder with various pathophysiological events that contribute to the pathogenesis and progression of the disease (Selkoe, 2011, Li and Selkoe, 2020). A key role for platelets in the progression of AD including CAA has been suggested by different groups (Stellos et al., 2010, Slachevsky et al., 2017, Donner et al., 2016, Jarre et al., 2014). Platelets store Aβ peptides and are able to release them upon platelet activation (Gowert et al., 2014, Chen et al., 1995, Li et al., 1998). The dysfunction of platelets is associated with AD and could be a contributor to AD pathology in the initiation and / or progression of the disease (Sevush et al., 1998, Johnston et al., 2008, Stellos et al., 2010, Slachevsky et al., 2017).

Since amyloid-antibody-therapies have been developed for AD patients, a comprehensive analysis of platelet activation and their contribution to the pathology of the disease is essential to develop effective blood screening tests Therefore, platelets might serve as important biomarkers for the diagnosis of AD.

In line with the results from the present study, several studies in the past showed unaltered platelet counts and size suggesting unaltered platelet turnover (Dos Santos and Pardi, 2020, Sevush et al., 1998). In contrast, Inestrosa and colleagues provide evidence for increased platelet counts (Inestrosa et al., 1993).

With regard to platelet activation, APP processing and protein expression in AD patients, controversial results have been shown by different clinical studies. Sevush and colleagues provide evidence for elevated CD62P (P-selectin) surface expression, platelet aggregate and platelet-leukocyte formation (Sevush et al., 1998). Järemo et al. showed that **s**oluble P-selectin was higher in the disease group, but not at the surface of platelets (Järemo et al., 2013). This study was based on 23 patients with moderate AD. In contrast, significantly higher baseline expression of activated integrin αIIbβ3 and P-selectin was observed in patients with AD with fast cognitive decline compared with AD patients with slow cognitive decline (Stellos et al., 2010). Here, we detected unaltered platelet activation under resting as well as under activated conditions when we examined platelets from patients at early stages of AD. However, when we separated the samples according to the sex of AD patients, we found differences between male and female patients. In specific, integrin aIIbb3 activation of female platelets following CRP and PMA stimulation was significantly reduced compared to platelets from males. Thus, it might be of great importance to analyze platelet activation of AD patients according to the sex of patients since we found no differences without separation of male and female platelets compared to healthy subjects. Furthermore, the differences in the activation profile of AD patients in different studies might be due to the stage of AD when the studies have been performed. In some clinical studies, no information has been given about the state of the disease. Moreover, it might be important to compare patients with different pace of cognitive decline as shown by Stellos and colleagues.

Further platelet parameters in AD patients have been elevated. This includes the number of coated platelets at early stages of AD that are decreased at late stages of the disease (Prodan et al., 2008). This further strengthens the need for the analysis of different stages of AD where platelet activation and related parameters could be different only because of the status of the disease. Interestingly, a study by Kumar and colleagues provided evidence for increased accumulation of neurotransmitter in platelets of female AD patients (Kumar et al., 1995). This further suggests that sex-specific analysis of platelet activation is essential to get valid results that might serve as reliable biomarker.

Different studies suggest to use alterations in platelets as prognostic biomarkers in AD. Stellos and colleagues found differences of platelet activation in patients with different velocity of cognitive decline (Stellos et al., 2010). Another studies from Ramos-Cejudo and colleagues suggests that increased platelet aggregation might have the potential as prognostic value for AD (Ramos-Cejudo et al., 2022). In addition, variations in platelet protein expression might serve as biomarker for AD. Yu and colleagues detected elevated protein expression of PHB, GPIBa and FINC while ADAM10 protein expression was decreased in platelets from AD patients (Yu et al., 2022, Yu et al., 2021). Differences in protein expression has been also observed by Fu and colleagues. They detected reduced ADAM10 expression as well but increased adenosine A2 receptor expression in platelets from AD patients (Fu et al., 2023). Other studies suggest to use the altered APP protein ratio in platelets from AD patients as potential biomarker (Di Luca et al., 1998, Zainaghi et al., 2012). Another promising candidate might be the mitochondrial function in platelets. Fisar and colleagues suggest that mitochondrial dysfunction could be a primary factor that contributes to the progression of AD (Fišar et al., 2019). This is in line with own experimental data showing that mitochondrial dysfunction contributes to platelet-mediated Aß aggregate formation at least in vitro (Donner et al., 2021). Here we show that determination of Aß at the platelet surface as detected by flow cytometry might be a promising candidate to serve as potential biomarker since we detected a strong increase in the binding capability of AD platelets to Aß. In addition, determination of platelet activation -at least in female platelets of AD patients-might be a valuable biomarker for the diagnosis of early AD in females. These parameters can be easily detected by flow cytometry. However, a more sophisticated approach might be the analysis of platelet morphology by TEM as we detected a significantly increase in the OCS of AD platelets. Beside the idea to serve as potential biomarker, the question about the role of an increased OCS arises and which consequences this phenotype might have for platelet-mediated AD progression. So far, there is only less known about the role of the OCS for platelet function. This internal membrane structure was identified 55 years ago and comprises a tunneling network of surface-connected channels (Selvadurai and Hamilton, 2018). However, it functions and how it is regulated is still not clear to date. In some human platelet disorders, structural abnormalities of the OCS have been detected but how they contribute to altered platelet function remains elusive.

Taken together, it will be of great importance to 1^st^ understand how platelets contribute to AD pathology in patients and 2^nd^ to design a panel of platelet parameters that might serve as biomarker for a first prognosis of AD because platelets can be easily accessed by blood withdrawal. These parameters might include platelet activation, the capability to bind Aß to the platelet surface and the dimension of the OCS by TEM.

## 6 Author contribution

ME and TS designed the study. LD performed experiments, JC and MK collected specimens. ME, TS and LD analyzed and interpreted data. ME wrote the manuscript with all authors providing feedback.

## 8 Acknowledgements

We thank Martina Spelleken for providing outstanding technical assistance.

## 10 Data Availability Statement

The original contributions presented in the study are included in the article, further inquiries can be directed to the corresponding author.

