## Supplemental Data for "Platelets from early-stage Alzheimer patients show enhanced amyloid binding, an elevated open canalicular system and sex-specific differences in their activation profile"

### *Supplementary Material*

#### 1 Supplementary Figures and Tables

##### 1.1 Supplementary Figures

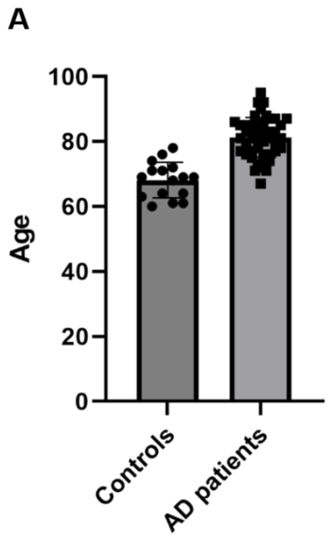


**Supplementary Figure 1. Overview of the age of AD patients and healthy subjects included in this study.** All subjects were >70 years of age. Bar graphs indicate mean values ± SEM. Statistical analyses were performed using an unpaired t-test. N = controls 17 and patients 46.

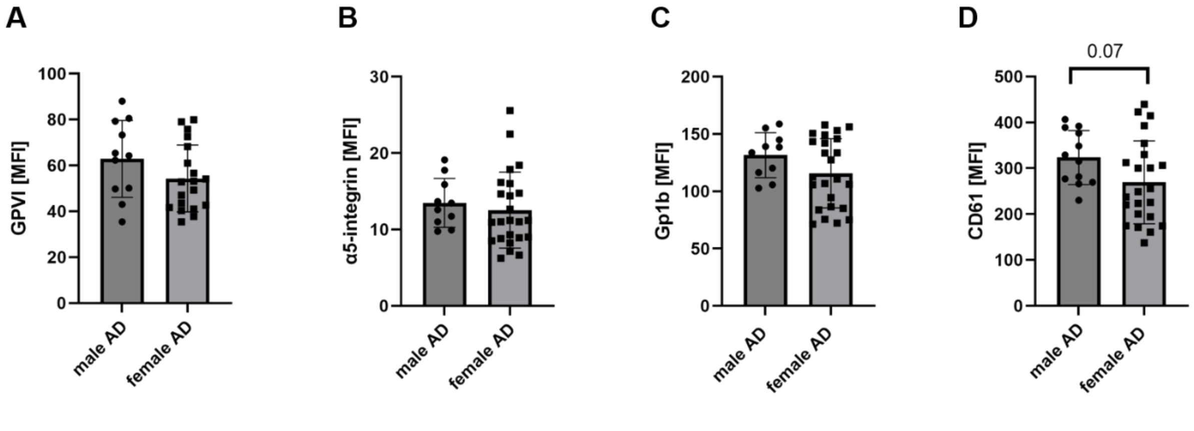


**Supplemental Figure 2. No differences in glycoprotein expression at the platelet surface between female and male AD patients**. Glycoprotein exposure at the platelet surface was determined by flow cytometry using different antibodies against (A) GPVI, (B) a5-integrin, (C) GPIb and (D) CD61 (subunit of integrin aIIbb3. Bar graphs indicate mean values ± SEM. Statistical analyses were performed using an unpaired t-test. N = 17 (controls), 35 (patients).

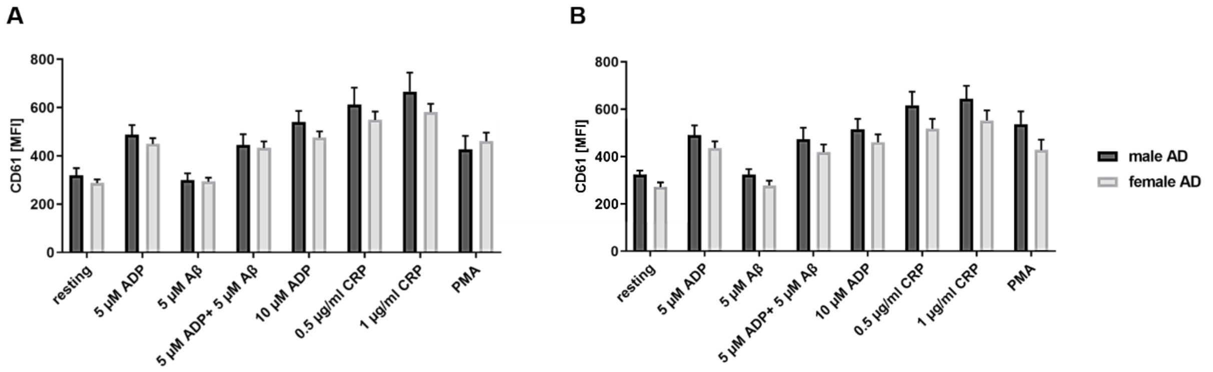


**Supplemental Figure 3.** No differences in the exposure of integrin aIIbb3 at the platelet surface between female and male AD patients. The exposure of integrin aIIbb3 under resting and under stimulating conditions was determined by flow cytometry using a CD61 antibody. Data are represented as MFI. Bar graphs indicate mean values ± SEM. Statistical analyses were performed using a multiple unpaired t-test. ADP, adenosinediphosphate, CRP, collagen-related peptide, Aß, amyloid-beta, PMA, Phorbol-myristate-acetate, MFI, mean fluorescence intensity. N = 17 (controls), 35 (patients).
